# Bamsnap-LRS: an automated batch visualization tool for long-read sequencing alignments

**DOI:** 10.64898/2026.06.21.733121

**Authors:** Wanyi Chen, Chentao Yang, Lingxin Qiu, Jiang Hu, Yang Zhou

## Abstract

**Summary:** Long-read sequencing (LRS) has become essential for genome assembly, structural variations (SVs) detection, haplotype phasing and transcript isoform characterization. However, these applications often require manual inspection of read alignment for validation. Existing visualization tools are either interactive genome browsers that are difficult to scale to large datasets or batch-oriented tools that are not optimized for the unique alignment patterns of long-read data. We developed Bamsnap-LRS, an automated command-line tool for high-throughput LRS alignment visualization. It supports long-read-specific features, phased SNP inspection, and publication-ready batch figure generation within a unified framework for genomic, transcriptomic, and haplotype-aware analyses.

**Availability and Implementation:** All codes and examples are freely available at https://github.com/comery/Bamsnap-LRS.

**Contact:** Chentao Yang and Yang Zhou.

**Supplementary information:** Supplementary Table 1 and Supplementary Figures 1-8 are available at *xxx* online.

## 1 Introduction

Long-read sequencing (LRS) technologies, represented by Pacific Biosciences (PacBio), Oxford Nanopore Technologies (ONT), and newly developed platforms such as CycloneSEQ and Qitan (Wang, *et al*., 2022; Zhang, *et al*., 2024; Zhang, *et al*., 2024), have transformed genomics (van Dijk, *et al*., 2023). By generating reads ranging from tens to hundreds of kilobases, LRS has greatly facilitated high-quality and even telomere-to-telomere (T2T) genome assemblies (Logsdon, *et al*., 2021; Mc Cartney, *et al*., 2022; Miga, *et al*., 2020; Nurk, *et al*., 2022; Yang, *et al*., 2023), while also demonstrating advantages in a broad range of applications, including structural variant (SV) detection, haplotype phasing and transcript isoform characterization (Monzo, *et al*., 2025; Olson, *et al*., 2023; Sedlazeck, *et al*., 2018). As LRS becomes increasingly adopted in population genomics, clinical genomics, and large-scale biodiversity initiatives, the volume of read-level evidence requiring interpretation has grown dramatically (Blaxter, *et al*., 2025; De Coster, *et al*., 2021; Liao, *et al*., 2023; Rhie, *et al*., 2021; Schreiber, *et al*., 2024).

**Figure 1.**
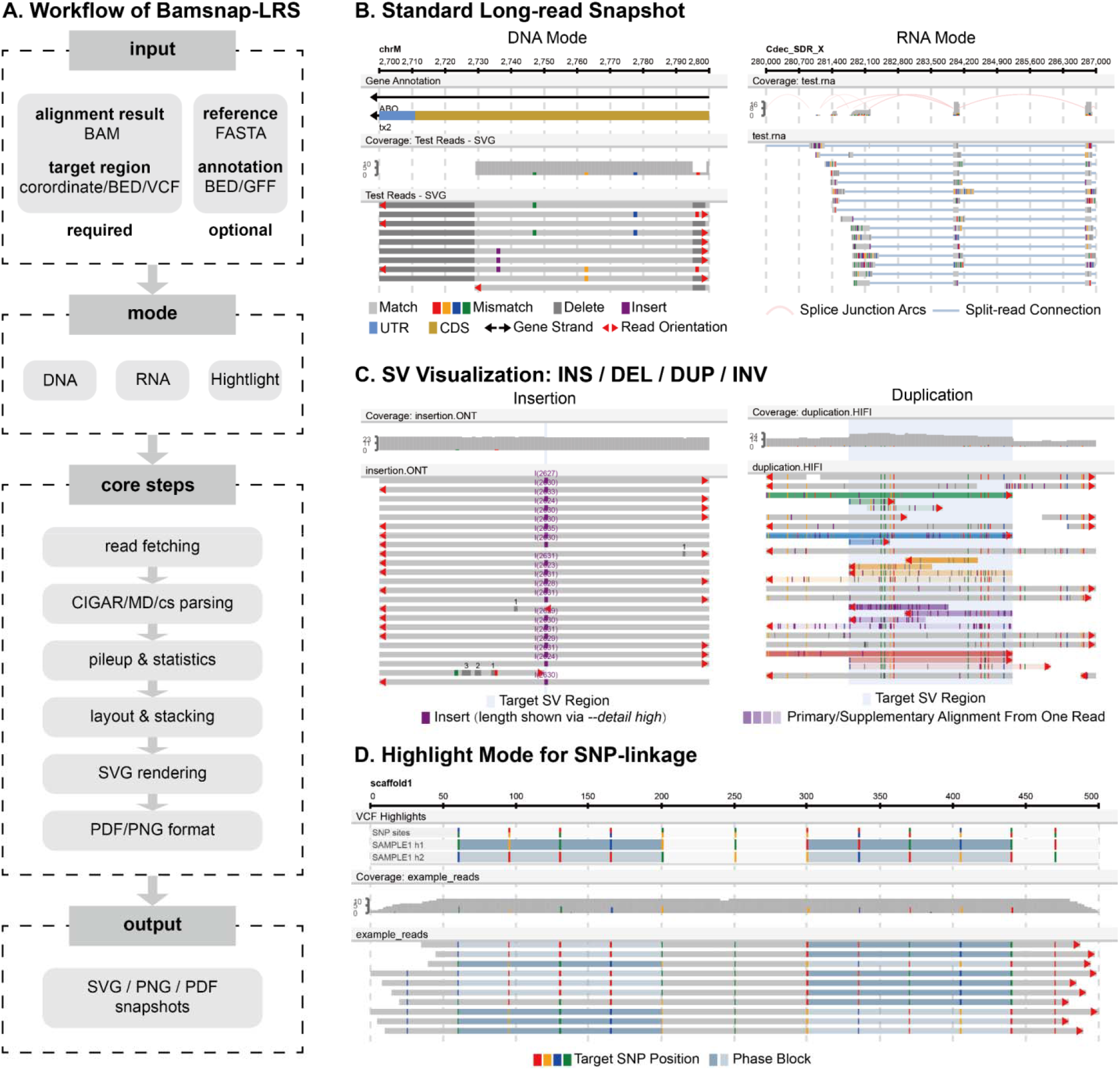
Overview and representative visualization modes of Bamsnap-LRS. (A) Bamsnap-LRS workflow from alignment input to final visualization output. (B) Representative long-read alignment snapshots produced in DNA and RNA modes, illustrating support for genomic and transcriptomic LRS data. (C) SV visualization examples. Bamsnap-LRS supports visualization of insertion, deletion, duplication, and inversion. Examples of an insertion and a duplication are shown. (D) Highlight mode for visualization of phased SNPs and read-level allele observations, facilitating SNP linkage and phasing assessment.

Despite the rapid advances in analytical methods, visualization remains a major bottleneck in LRS data analysis. Many important biological conclusions—including assembly validation, SV discovery, haplotype phasing and transcript characterization—ultimately depend on direct inspection of read alignment. For example, long reads and specialized SV callers, including Sniffles2 (Smolka, *et al*., 2024), CuteSV (Jiang, *et al*., 2020) and SVision-Pro (Wang, *et al*., 2025), have substantially improved SV detection, particularly within repetitive regions (Sedlazeck, *et al*., 2018). However, complex genomic regions and alignment ambiguities can still generate a considerable number of false positives that need to be manually reviewed (Cretu Stancu, *et al*., 2017; Qin and Li, 2025; Zook, *et al*., 2020). Similarly, in haplotype phasing, tools such as WhatsHap (Martin, *et al*., 2016) and HapCUT2 (Edge, *et al*., 2017) can accurately reconstruct haplotypes from long reads, but switch errors, fragmented phase blocks, and ambiguous haplotype assignments can arise from sequencing errors, insufficient read support, or complex genomic regions (Sakamoto, *et al*., 2022) and often require careful validation. Long-read RNA sequencing further increases the demand for visualization by enabling direct observation of transcript structures and alternative splicing events. Collectively, these applications highlight the need for scalable visualization tools that can efficiently present diverse forms of read-level evidence for validation and interpretation of LRS results.

The most widely used visualization tools are the Integrative Genomics Viewer (IGV) (Robinson, *et al*., 2011) and JBrowse (Diesh, *et al*., 2023). While highly flexible, IGV and JBrowse are primarily designed for interactive graphical exploration. When thousands of variants need to be evaluated across hundreds of samples, manual navigation becomes labor-intensive. Several tools have been developed to address this limitation. For example, Bamsnap (Kwon, *et al*., 2021) enables batch visualization but was originally designed for short-read data, thus is not optimized for the distinctive alignment patterns of long reads, including extensive soft clipping, large indels, split alignments, and chimeric mappings. One recently published software SVHawkeye (Xiao, *et al*., 2024), particularly designed for long reads, resolved many of the above batch visualization issues for LRS alignment and introduced the function of SV genotyping. However, its main design focuses on batch visualization of SVs and is less suitable for more general visualization needs (e.g. customized annotation track) and other LRS applications, such as assembly validation, base-level variant inspection and haplotype-resolved SNP linkage visualization (**Supplementary Figs. 1-3**).

To address these challenges, we developed Bamsnap-LRS, a command-line visualization tool designed for generating publication-ready snapshots of LRS alignments. Building upon the batch-processing philosophy of the original Bamsnap, Bamsnap-LRS is additionally optimized for long-read sequencing data, with supporting visualization of structural variants, small variants, spliced transcript alignments, and haplotype phasing evidence. By integrating diverse long-read sequencing applications within a single reproducible workflow, Bamsnap-LRS enables a scalable solution for visualization, validation and interpretation of LRS data.

## 2 Software description

### 2.1 Overview

Bamsnap-LRS is an automated command-line visualization tool. Unlike interactive GUI-based tools, Bamsnap-LRS focuses on batch visualization of read alignments of predefined genomic regions, facilitating variant review, comparative visualization across samples, and publication-ready figures. The overall workflow of Bamsnap-LRS is shown in **Figure 1A**.

Bamsnap-LRS provides three visualization modes: DNA, RNA, and Highlight. DNA mode displays standard long-read alignments, coverage, and SV evidence within genomic regions. RNA mode visualizes spliced alignments and transcript structures from long-read transcriptomic data. Highlight mode integrates SNPs from the input VCF with LRS alignment to facilitate the inspection of SNP linkage and haplotype consistency.

Bamsnap-LRS requires one or more BAM files and target regions as input. When multiple BAM files are provided, alignments from each file are displayed in separate tracks. Target regions can be specified directly as a string, or supplied in BED or VCF for batch processing. BED files are parsed as genomic intervals, whereas VCF files are interpreted according to variant types and related information. During batch visualization, Bamsnap-LRS can automatically infer the plotting window around each target region. Insertions and small variants are displayed using relatively compact windows, and large structural variants are visualized using broader windows based on their estimated lengths. The software also provides parameters that allow users to customize the padding region on both sides of the target interval. Optional inputs include a reference genome FASTA file and gene annotation files in BED, GFF, or GTF format, allowing genomic features to be displayed alongside the alignment tracks. Highlight mode additionally requires a VCF file containing the target SNPs.

Bamsnap-LRS generates static output figures in SVG, PNG, or PDF format. SVG serves as the primary rendering format, and PNG/PDF outputs are generated from the same SVG-based rendering workflow to ensure consistent appearance across different output formats.

### 2.2 Implementation and Visualization Workflow

Bamsnap-LRS is implemented in Python 3. Its main workflow includes read fetching, alignment-tag parsing, pileup calculation, layout assignment, and snapshot rendering (**Figure 1A**). After determining the target regions, Bamsnap-LRS uses Pysam (https://github.com/pysam-developers/pysam) to fetch alignments from BAM files and convert them into standardized read objects. Each read object records the read name, genomic coordinate, strand, mapping quality, alignment status (primary or supplementary) and alignment-derived features. These features, including matches, mismatches, insertions, deletions, skipping and clipping, are mainly extracted from CIGAR strings and can be further refined using MD tag, cs tag, and reference sequence information.

After read parsing, Bamsnap-LRS calculates coverage, then arranges visual layouts for tracks such as coverage, annotation and read alignment. For long-read alignments, primary and supplementary alignments from the same read are displayed in a compact arrangement with colored fills, thereby preserving relevant alignment information while avoiding excessive visual clutter. In RNA mode, exon-spanning alignment segments are connected to represent spliced alignment structures. In Highlight mode, reads can be filtered and sorted according to whether they overlap target VCF sites and according to their observed haplotype signatures. Finally, the main rendering engine uses xml.etree.ElementTree and xml.dom.minidom to generate SVG snapshots, while PNG and PDF outputs are generated through CairoSVG (https://github.com/Kozea/CairoSVG) conversion.

## 3 Examples of Usage

### 3.1 Case 1: Standard Long-read Snapshot for DNA and RNA Alignments

As a visualization tool for long-read alignments, one of the primary functions of Bamsnap-LRS is to generate clear snapshots of basic alignment for both DNA and RNA reads (**Figure 1B**). In the DNA mode, the tool displays coordinate axes, coverage tracks, read pileups with mismatches, indels and read orientation. In RNA mode, Bamsnap-LRS renders transcript reads spanning large genomic intervals and multiple exons. Splice junction arcs summarize skipped genomic regions that usually correspond to introns in spliced transcripts, allowing users to rapidly inspect exon connectivity and potential alternative splicing patterns.

### 3.2 Case 2: SV Visualization

Bamsnap-LRS can also be used to visualize SVs (**Figure 1C**; more examples at https://github.com/comery/Bamsnap-LRS/tree/main/example/SV_show). As shown in **Figure 1C**, Bamsnap-LRS automatically extends the surrounding genomic interval for plotting, placing the target SV region near the center of the final snapshot. The target SV region is highlighted with a light shaded background. Insertions and deletions are shown as purple insertion blocks and dark gray blocks, respectively. For more complex SVs such as duplications and inversions, Bamsnap-LRS displays both primary and supplementary alignments from the same read as a compact group using a shared color scheme with an opacity gradient and arranging the alignments according to the order along the original read. This representation preserves the continuity of split-read alignments and facilitates interpretation of complex rearrangements. In duplication, primary and supplementary alignments typically share the same orientation, and the duplication size can be assessed from the genomic interval between the end of one alignment segment and the start of its adjacent segment on the same read. In inversion, alignment segments have opposite orientations, and the inversion size corresponds to the genomic span of the inverted segment between its flanking alignments.

### 3.3 Case 3: Highlight Mode Visualization

Highlight mode is used to visualize long-read alignments associated with SNPs from a VCF for assessing SNP linkage (**Figure 1D**). This mode contains a SNP track above the read pileup. Within the alignments, only bases corresponding to the target SNP sites in VCF are highlighted, while non-target mismatches and indels are suppressed to reduce visual noise. Reads are automatically grouped according to their observed SNP combination, even without phasing information. When phasing information is provided in VCF, SNPs from the same phase block are shown with a shared background color. Bamsnap-LRS further compares the allele combination observed on each read with the phased SNPs in the VCF, and labels supporting read segments with the corresponding phase block colors. This representation enables intuitive inspection of haplotype consistency, phasing accuracy, and potential linkage relationships between neighboring phase blocks.

## 4 Discussion

In summary, Bamsnap-LRS provides a unified command-line framework for generating snapshots of long-read sequencing alignments. By integrating multiple visualization modes, Bamsnap-LRS reduces the need for users to switch among multiple tools across different long-read sequencing applications. We summarize features of Bamsnap-LRS and some other visualization tools (**Supplementary Table 1**). Compared with interactive genome browsers, Bamsnap-LRS is not intended to replace manual inspection in IGV or JBrowse, but rather to complement them by providing automatic functionality, allowing researchers to choose complementary strategies according to their specific needs. Compared with existing batch visualization tools, Bamsnap-LRS places greater emphasis on the unified representation of long-read-specific features and diverse application scenarios (**Supplementary Figs. 1-8**). In addition, the Highlight mode provides a visual strategy for the manual inspection of SNP phasing results.

Bamsnap-LRS still faces some challenges. Because translocation usually involves inter-chromosomal relationships that are difficult to represent within a single linear genomic snapshot, such events are not yet fully supported. In addition, for extremely high-depth datasets or very large genomic regions, appropriate read filtering, downsampling, or region selection may still be required. Future development will focus on extending support for more complex rearrangements, integrating additional long-read signals such as methylation, improving performance for large-scale datasets, and providing more flexible visualization options for diverse sequencing applications.

## Author contributions

Conceptualization: C.Y., Y.Z.; Supervision: C.Y., Y.Z.; Data Curation: W.C., Y.Z., L.Q.; Software: W.C., C.Y.; Funding Acquisition: C.Y., Y.Z.; Writing—original draft: W.C., Y.Z.; Writing—review & editing: W.C., Y.Z., C.Y., J.H.

## Funding

This work was supported by grants from the National Key R&D Program of China (2025YFC3410300) to C.Y. and Shenzhen Medical Research Fund (D2501006) to Y.Z..

## Acknowledgement

We thank Yudian Peng and Dongming Fang for testing the software. We would like to thank DCS Cloud (https://cloud.stomics.tech/) for providing the computational resources and software support necessary for this study.

## Notes

### Competing Interest Statement

The authors have declared no competing interest.

