## Supplementary Table 1, Supplementary Figs. for "Bamsnap-LRS: an automated batch visualization tool for long-read sequencing alignments"

### **Supplementary Table 1: Summary of Bamsnap-LRS and some other visualization tools**

| Items | IGV | Bamsnap | SVhawkeye | Bamsnap-LRS |
| --- | --- | --- | --- | --- |
| LRS supported | √ | × | √ | √ |
| Requirements | Java | python (Pysam, Pillow, pyfaidx, pytabix) | python (Pysam), R (getopt, data.table, RColorBrewer) | python (Pysam, CairoSVG) |
| Required Input | Reference (FASTA)  Alignment file (BAM/CRAM) | Reference (FASTA)  Alignment file (BAM/CRAM)  Genomic positon (coordinate/VCF/BED) | Alignment file (BAM)  Genomic positon (VCF/BED) | Alignment file (BAM/CRAM)  Genomic positon (coordinate/VCF/BED) |
| Optional Input | Gene Annotation (BED/GFF/GTF)  Variant Records (VCF)  Custom Interval Statistics (BED/BigWig) | - | Reference (FASTA) | Reference (FASTA)  Gene Annotation (BED/GFF/GTF)  Variant records (VCF) |
| Output Format | PNG, SVG | PNG, JPG | PNG, PDF | SVG, PNG, PDF |
| Batch Supported | × | √ | √ | √ |
| Support Type | Standard alignment display | Standard alignment display | SV, RNA splicing | Standard alignment display, SV, RNA splicing |
| Additional Features | - | - | SV genotyping, CNV depth | SNP highlight |
| Speed (sec/SV)* | - | - | 1.47 | 0.83 |

*Since Bamsnap-lrs is a single-threaded tool, SVhawkeye also uses a single thread when testing speed.

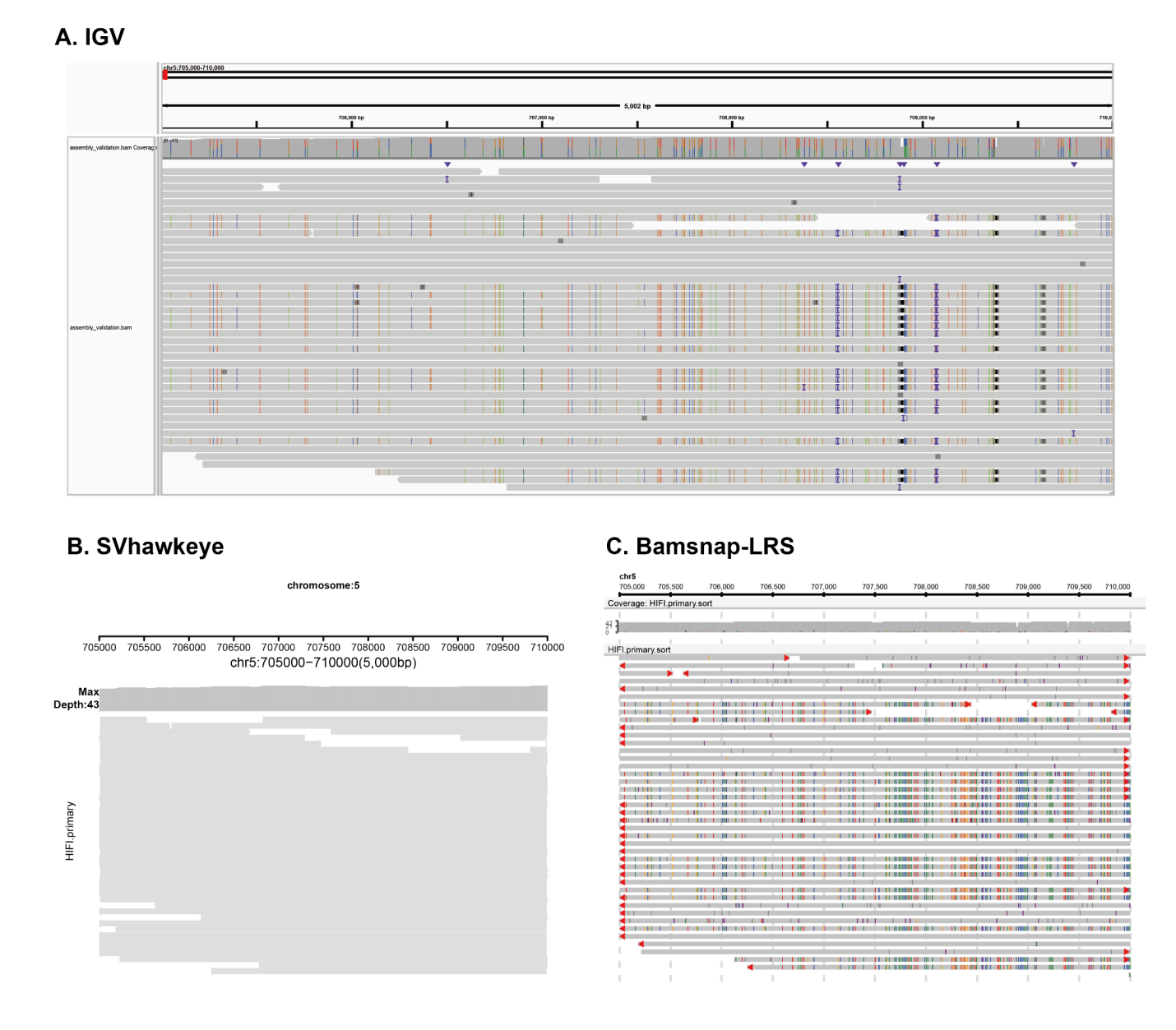

### **Supplementary Figure 1. Visualization of long-read alignments of a rare collapsed region in CHM13** (Mc Cartney, *et al.,* 2022) **using IGV (A), SVHawkeye (B), and Bamsnap-LRS (C).**

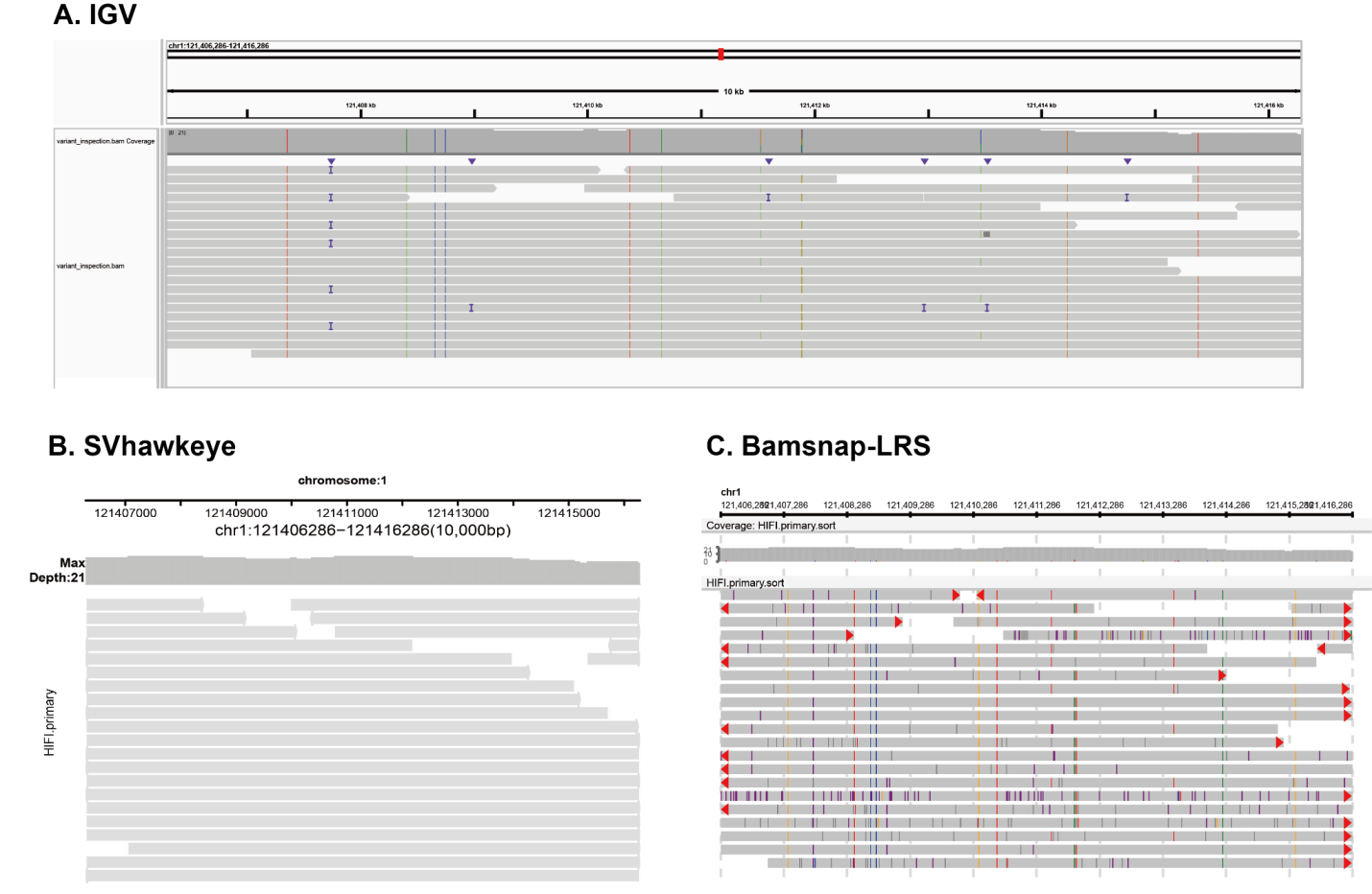

### **Supplementary Figure 2. Visualization of long-read alignments for base-level variant inspection in a centromere region (CHM13 chr1:121406286-121416286) using IGV (A), SVHawkeye (B), and Bamsnap-LRS (C).**

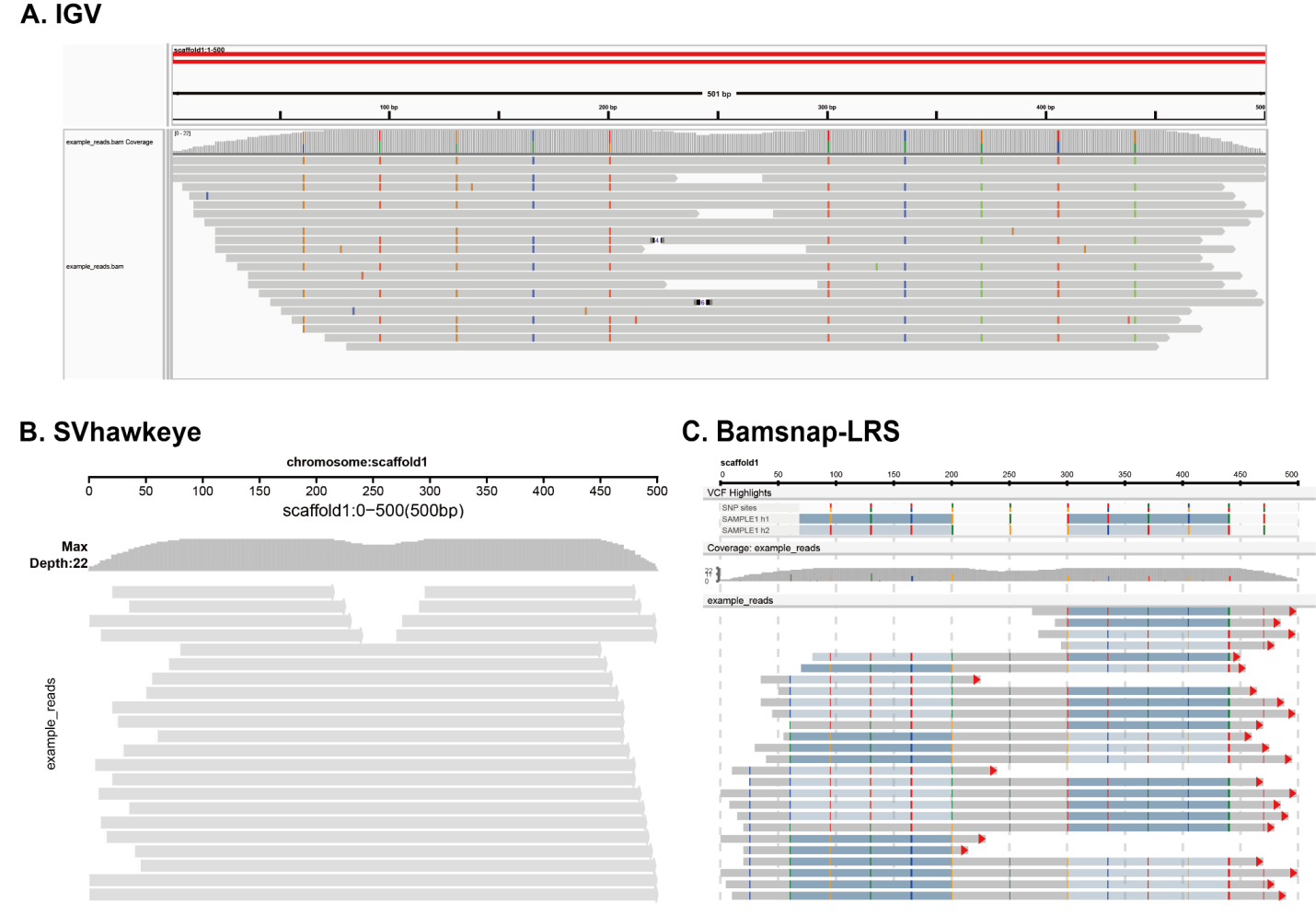

### **Supplementary Figure 3. Haplotype-resolved SNP linkage visualization.** In Bamsnap-LRS, supporting read segments are colored according to the corresponding haplotype in VCF without haplo-tagging.

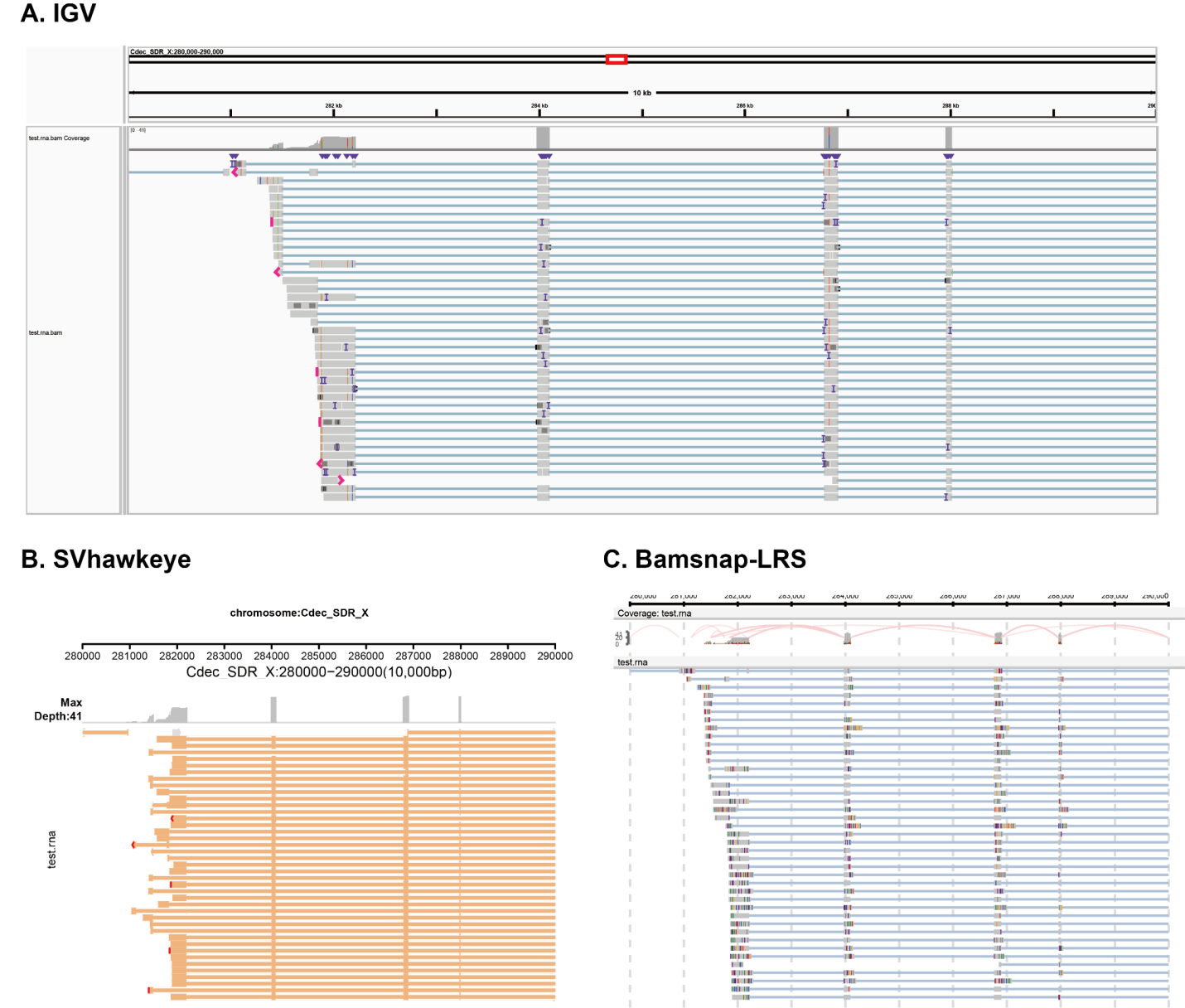

### **Supplementary Figure 4. Transcript isoform visualization using IGV (A), SVHawkeye (B), and Bamsnap-LRS (C).**

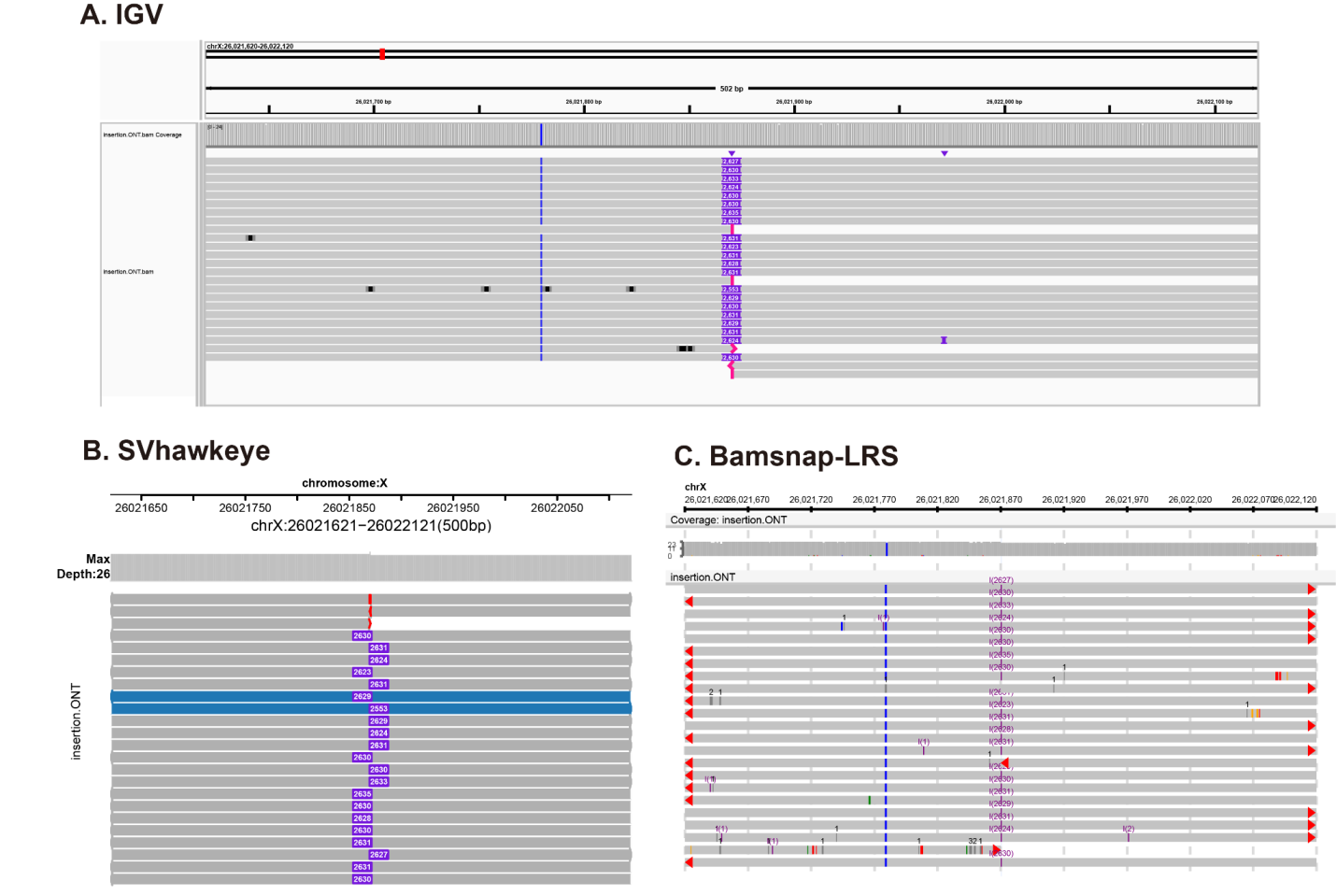

### **Supplementary Figure 5. Insertion visualization using IGV (A), SVHawkeye (B), and Bamsnap-LRS (C).**

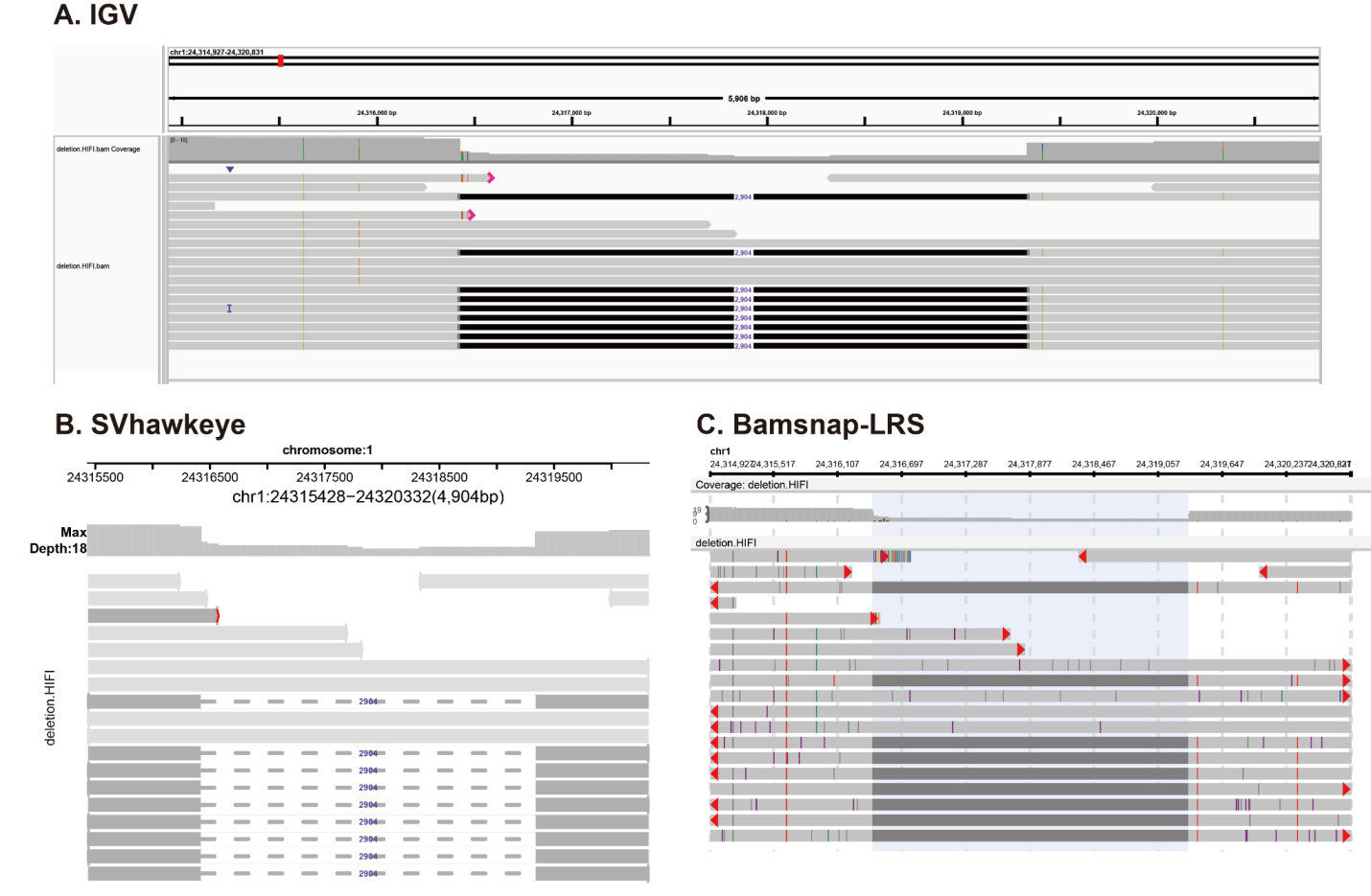

### **Supplementary Figure 6. Deletion visualization using IGV (A), SVHawkeye (B), and Bamsnap-LRS (C).**

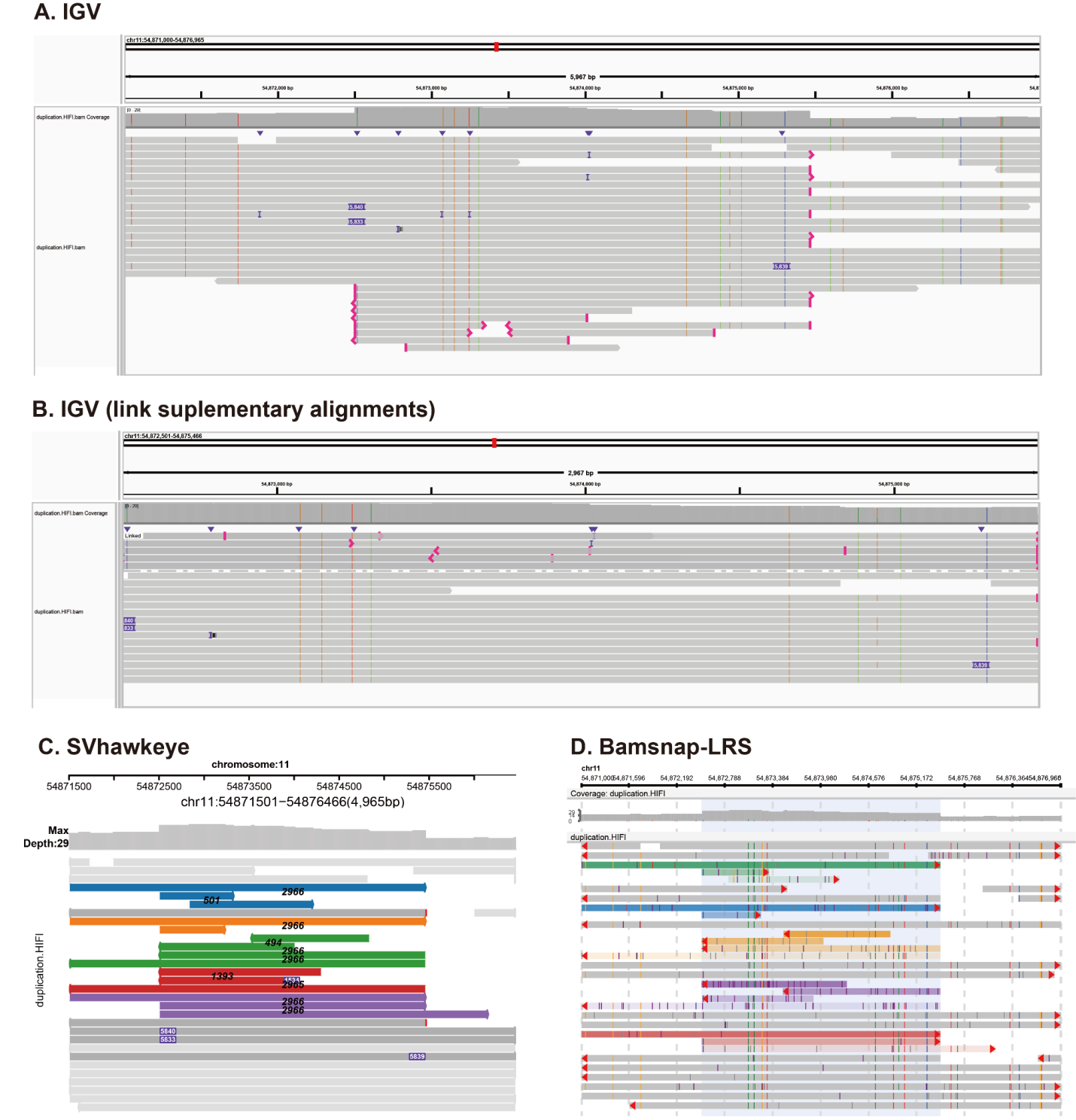

### **Supplementary Figure 7. Duplication visualization using IGV (A), IGV with “link supplementary alignments" option (B), SVHawkeye (C), and Bamsnap-LRS (D).**

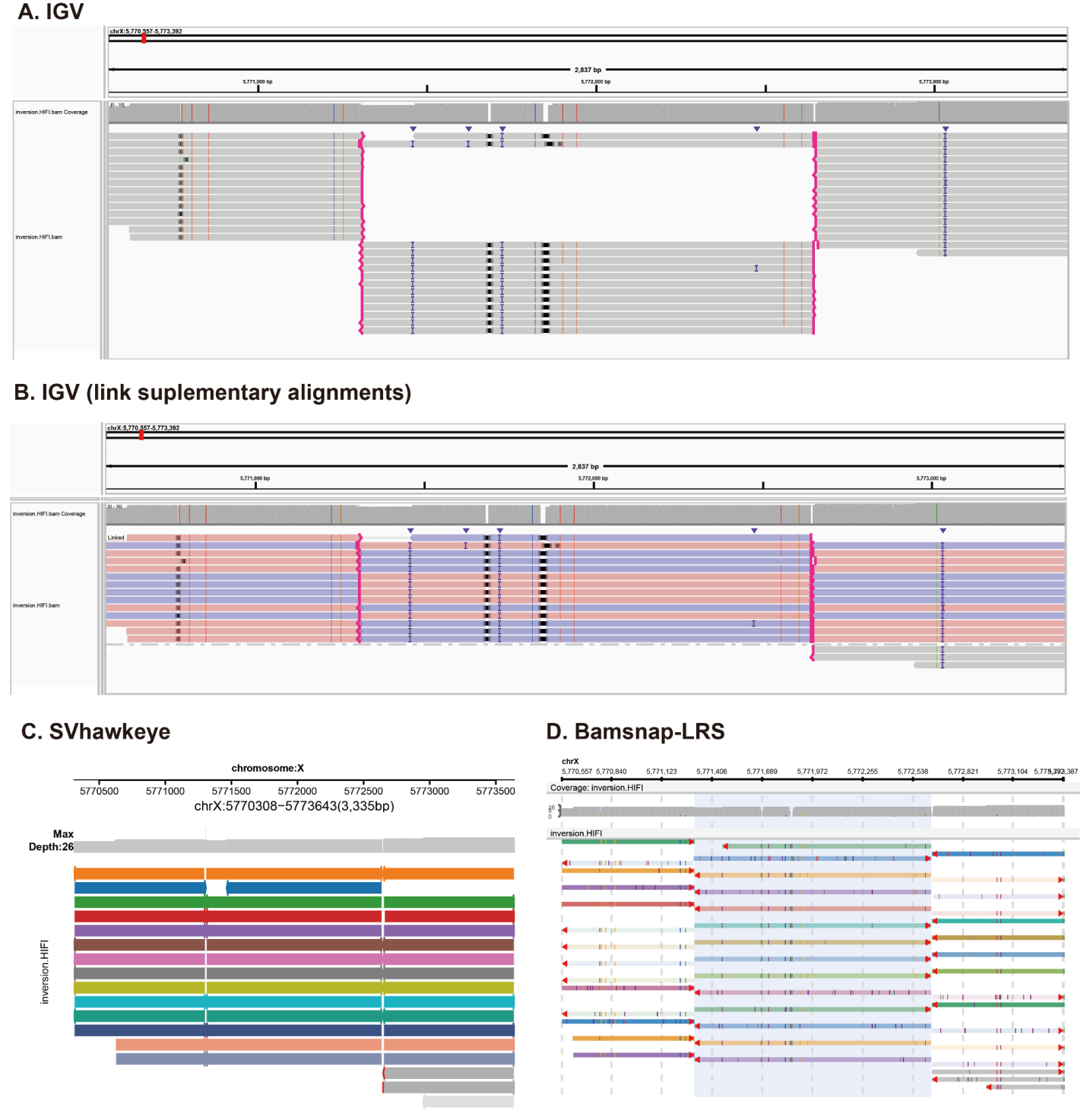

### **Supplementary Figure 8. Inversion visualization using IGV (A), IGV with “link supplementary alignments" option (B), SVHawkeye (C), and Bamsnap-LRS (D).**
